# KSTITCH links cellular morphology and gene expression in spatial transcriptomics

**DOI:** 10.64898/2026.06.07.730714

**Authors:** Shashwat Kumar, Yingxiao Shi, Tuulia Vallius, Chi-Ping Day, P.-A. Absil, Anuj Srivastava, Sridhar Hannenhalli, Vishaka Gopalan

## Abstract

In situ spatial (ISS) sequencing can uncover co-variation between cellular morphology and gene expression in vivo. However, a principled and interpretable mathematical representation of morphology has not yet been applied in this context. In particular, current deep learning-based representations of cell images confound a cell’s shape with its size. We present an interpretable representation of cellular boundary contours, based on tangent principal component analysis (TPCA) in a Kendall shape manifold, that captures size-independent contour shape features. This approach successfully recovers shape-perturbing genes in an RNAi screen than a previous metric geometry-based approach. We build on TPCA to develop KSTITCH (Kendall Shape–TranscriptomIc Correlation and Harmonization), an approach to reveal covariation between cell morphology with gene expression in ISS datasets. In a Xenium dataset, KSTITCH recovers known morphology-transcriptomic relationships in keratinocytes, macrophages and endothelial cells. Across samples in a melanoma CosMx dataset, KSTITCH reproducibly associates elongated and triangular fibroblasts with proximity to malignant cells and myofibroblast-like transcriptional program. Finally, KSTITCH independently recovers a known link between mesenchymal-like malignant cell states and increased cell area in two melanoma cohorts. KSTITCH can thus yield interpretable morphology-transcriptome relationships across cell types, patients, and spatial transcriptomics platforms. KSTITCH is available at https://github.com/vishakagopalan/kstitch.

## Introduction

A cell’s morphology and its transcriptional state are bidirectionally linked, where perturbing morphology causes transcriptional change[1, 2], and transcriptional perturbations result in morphological changes [3, 4, 5]. Pioneering studies jointly investigating the morphology of cancer cell lines with their bulk RNA-seq profiles found links between a cell line’s gross nuclear morphology and its invasiveness [6] and metastatic potential [7] in addition to correlations between diverse signaling pathways and cellular contour shape and size [8, 9, 10]. Spatial in situ sequencing (ISS) transcriptomics platforms such as CosMx and Xenium that provide paired cellular/nuclear morphology and transcriptomic data from tumors at single-cell resolution now allow us to extend the previous investigations, largely performed in vitro, to in vivo settings. Here we present an approach based on analyzing cell contours using Kendall’s shape analysis [11, 12, 13] to identify interpretable axes along which transcriptional state and cell shape covary within a tissue.

Existing computational pipelines vary both in their approaches to model contours and in their downstream analysis goals. MUSE [14] is built for clustering cells based on both their morphology and transcriptomic readouts but is not designed to infer which genes’ expression co-vary with specific, interpretable modes of morphological variation. MorphLink [15] links spatial gene expression variation to morphological variation in nuclei and other tissue structures using interpretable summary descriptors such as circularity, eccentricity, solidity, area, and measures of tissue organization. However, MorphLink does not link transcriptional programs in cell states to variation in their morphology. Thus, neither MUSE nor MorphLink is directly comparable to our tool, which aims to specifically learn interpretable, yet not explicitly pre-defined, cellular or nuclear contour features and reveal gene expression programs associated with these learned features.

CAJAL [16], in contrast, directly compares contours by computing Gromov-Wasserstein pairwise distances between them. However, these pairwise distances do not provide an explicit representation of contour variation, decompose that variation into interpretable modes, or yield a generative model of cell shape. More importantly, CAJAL distances do not distinguish variation in cell size from variation in shape, which can reflect distinct biological processes [17, 18]. This conflation is particularly problematic in ISS datasets, where differences in cellular or nuclear size can also arise from technical effects, including field-of-view edge effects in CosMx datasets [19] and differences in image magnification across slides.

We resolve this shortcoming by computing size-independent representations of cellular and nuclear contours from ISS data on a Kendall shape manifold where a cell’s shape is defined as the geometrical information of a contour that remains after the contributions of its size, location and orientation are marginalized. Further, we perform tangent principal component analysis (TPCA) of contour data on the Kendall shape manifold to discover dominant modes of shape variation in a population. Each shape PC is interpretable by visual inspection or by standard statistical tests, e.g., by correlating them with such pre-determined morphological features as eccentricity and circularity. Euclidean distances in TPCA space are designed to approximate geodesic distances between shapes on the Kendall manifold. This is in contrast to earlier pipelines that perform PCA on contour coordinates without an explicit manifold representation step [20, 21, 22].

Additionally, we study gene expression factors in ISS data and represent them as Euclidean vectors using non-negative matrix factorization (NMF) of read counts. Our main contribution is a novel pipeline, termed KSTITCH (Kendall Shape–TranscriptomIc Correlation and Harmonization), which uses Canonical Correlation Analysis (CCA) to find combinations of gene expression factors that highly correlate with combinations of cellular or nuclear shape PCs and size. Importantly, we model all technical covariates of size, shape PC, and factor variation within and between ISS slides prior to the CCA step. We first validate TPCA’s ability to detect abnormal nuclear shape in a published RNAi screen of shape-perturbing genes where TPCA outperformed CAJAL in terms of detecting perturbed nuclei without being confounded by nuclear size. We then demonstrate KSTITCH’s ability to reproducibly discover known patterns of shape-gene covariation among multiple cell types in publicly available CosMx- and Xenium-derived melanoma datasets. In particular, KSTITCH recovered known morphology-gene expression relationships in keratinocytes, endothelial cells, and macrophages in a Xenium Prime dataset. KSTITCH also reproducibly recovered shape changes associated with fibroblast activation in proximity to tumors as well as a known link between melanoma de-differentiation and cell size. These results demonstrate the wide applicability of KSTITCH to multiple cell types and spatial transcriptomics platforms.

## Results

### Tangent Principal Component Analysis generates shape representations that can help detect nuclei with perturbed shapes

A two-dimensional contour is represented by a 2 × *L* matrix of coordinates, where L denotes the number of landmarks on the object contour, each having a (*x, y*) coordinate. Shape is the property of an object that is retained after we have removed the influence of its position, size, and orientation [12, 11, 13]. In addition to these transformations, since the choice of the first landmark on a closed contour is arbitrary, one also needs to match the starting landmarks between a pair of shapes in order to quantify their shape differences. Once we account for these quantities (translation, scaling, rotation, and starting landmark) in a shape’s landmark matrix, the resulting representation is no longer in a traditional Euclidean space, but rather in a nonlinear manifold where one uses geodesic distances – lengths of shortest paths on that manifold – to quantify shape differences. However, these distance computations can be cumbersome, especially for large datasets. To gain efficiency, one uses an approximation called tangent principal component analysis (TPCA),where each shape is represented in a parsimonious fashion as a (Euclidean) principal component vector that captures much of the variation of the data.

Figure 1a is an overview of TPCA (see Methods for details). Since cellular and nuclear segmentation methods can generate contours with varying numbers of landmarks for each shape, we first interpolate each shape and uniformly sample L landmarks from it. In Step 2, we mean-center all shapes and normalize their areas to one. In Step 3, we simultaneously compute their sample mean shape and register individual shapes to the mean shape using an iterative scheme. The latter implies that we eliminate differences in orientation and the position of the first landmark between shapes. Each iteration is composed of two steps: (a) estimating the mean of all shapes in the dataset, and (b) rotating each individual shape and repositioning the first landmark on it using the mean shape as a reference (see Methods). Upon convergence, the mean shape is computed, and a principal component (PC) embedding of all shapes is determined in Step 4. Though inter-shape distances based on TPCA represent a Euclidean approximation of the geodesic distances, we verify these approximations in our applications. Much like classical PCA, pairs of PCs represent orthogonal modes of variation.

**Figure 1.**
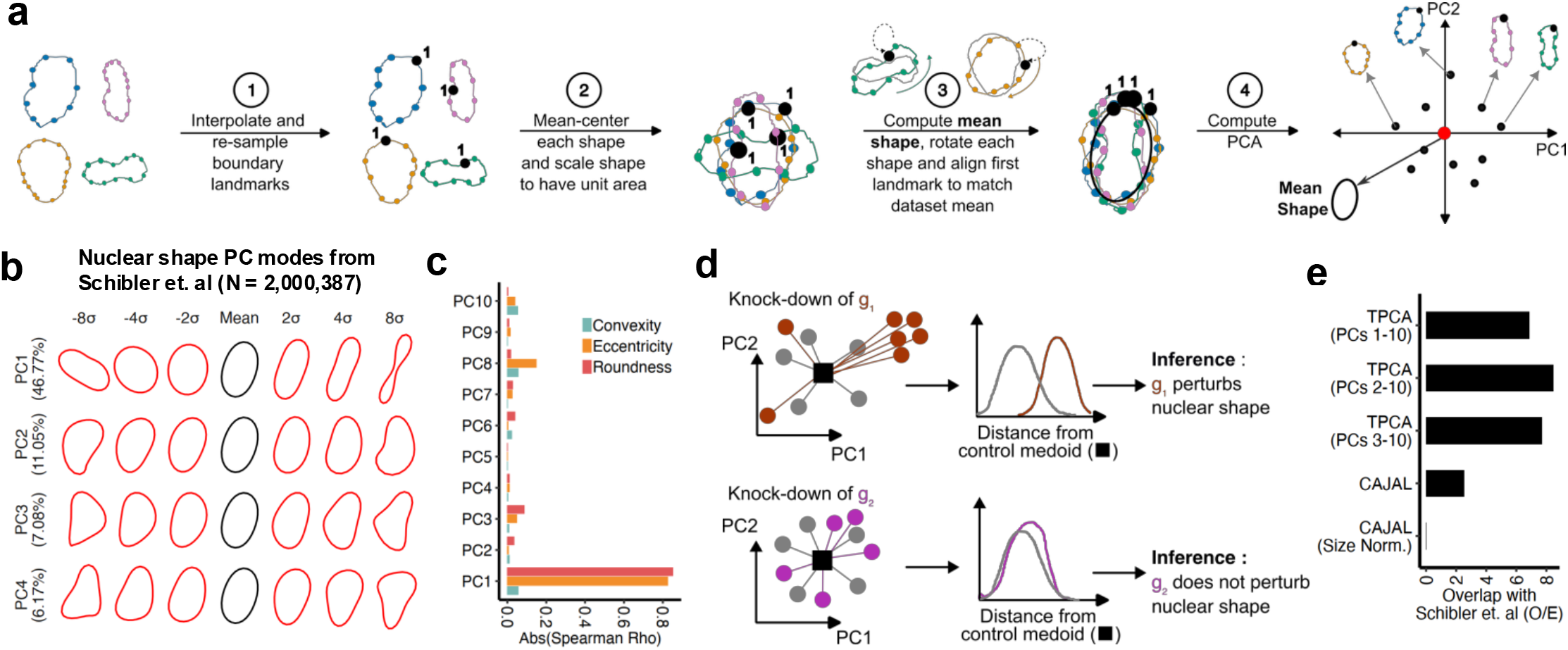
(a) Overview of tangent principal component analysis (TPCA) algorithm. A fixed number of landmarks are resampled from each shape’s boundary using linear spline interpolation. The first landmark is indicated with a large black circle. Shapes are translated and scaled such that landmark coordinates of each shape sum to zero and each shape’s landmark matrix has a Frobenius norm of one. The Frechét mean of all shapes is then computed, and each shape is rotated (and its first landmark moved) to align with the mean. Finally, shape matrices are represented as vectors in a tangent space around the mean shape and a PCA vector of each shape computed. (b) Estimate of the mean shape of all nuclei in Schibler et. al data (black), with shapes in red sampled between +8 and −8 standard deviations from the mean shape along each PC direction. Variance explained by each PC is shown in brackets. (c) Correlation of each shape PC derived from nuclei in Schibler et. al data with the convexity, eccentricity and roundness (or circularity) of nuclei. (d) Scheme used to detect nuclear shape-modifying genes using CAJAL or TPCA. The medoid of control nuclei (black square) is computed, after which the distances of nuclei perturbed by a given gene is computed. A gene (in brown) whose knockdown increases the distance of nuclei from the control medoid (black square) is labelled as shape-modifying while a gene (in purple) that does not increase distance is not considered to modify nuclear shape. (e) Observed over expected ratio of overlap between shape-modifying genes detected by TPCA and CAJAL with the gold-standard set defined by Schibler et. al.

We first demonstrate the utility of the TPCA approach in identifying genes that control nuclear shape on a published nuclear morphology RNAi screen conducted in immortalized fibroblasts by Schibler et al [23]. In that study, the authors tested the effect of knocking down each of 861 genes using three siRNA probes each to identify 34 genes (based on their effect on nuclear eccentricity and circularity); we consider this a ground truth gene set. We re-processed raw images from this data using the CellPose algorithm to segment raw images of nuclei based on DAPI intensity signals and derived contours of 2 million nuclei (see Methods) from this screen. We uniformly sampled fifty landmarks along each nuclear contour and ran the TPCA procedure on all nuclei. The first ten PCs captured 90% of the variance (Supp. Fig. 1a) and were retained for downstream analyses. TPCA embeddings were robust to batch effects, i.e., nuclei perturbed by the same siRNA probe across biological replicates had similar embeddings (Supp. Fig. 1b, Methods). TPCA yields a generative and invertible representation of shapes in a dataset such that a shape can be represented by a PC vector. The mean shape of all nuclei (black, Fig 1b) is found to be elliptical, and the eigenvectors in TPCA space can be used to recreate a nuclear shape from its PC vector. Nuclear shapes in red (Fig. 1b) are computed from PC eigenvectors up to +/- 8 standard deviations respectively from the mean along each tangent principal direction. We compared popular descriptors of shapes such as eccentricity, roundness and convexity (ratio of the convex hull perimeter to the observed shape perimeter) with each PC to interpret their contributions to shape variation (Fig. 1c). Overall, PC1 captured 46% of shape variation and was most strongly correlated with eccentricity and roundness. Other PCs were weakly correlated with these shape descriptors. Importantly, distances between nuclei computed based on the all PCs and the first ten shape PCs (Supp. Fig 1c) was tightly correlated with geodesic distances between them (Rho = 0.981 for first ten PCs) which justified the use of TPCA-based distances in our applications.

Next, we assessed the extent to which our approach can identify shape genes that were experimentally identified by Schibler et al. We reasoned that the shapes of nuclei perturbed by the control siRNA probes represent background shape variation and therefore we computed as a reference null distribution, the distances of all control nuclei from their medoid (Fig. 1d). For each perturbation, we checked if perturbed nuclei were more distant from the control medoid than the null distribution using a one-sided Wilcoxon test. We considered a gene to be a shape gene if at least two out of three targeting siRNAs had a false-discovery rate (FDR) less than 0.1. Based on the first ten PCs, TPCA labelled 21 genes as shape genes of which seven overlapped the ground truth (Odds Ratio = 6.84, auPRC enrichment over baseline = 3.10, Fig. 1e, Supp. Fig. 1d). Since PC1 correlated with roundness and eccentricity, which were the features used by Schibler et al. to identify shape genes, we repeated our distance calculation without PC1 and PC2 to test if later PCs could still identify shape genes. Omitting PC1 yielded 17 candidate shape genes, while omitting the first two PCs yielded 16 candidate genes, both of which represented an odds ratio enrichment of 8.45 and 7.69, respectively. We repeated the procedure in Fig 1d using CAJAL[16] and found that it recovered all ground truth genes but labelled 277 genes as candidate shape genes (Fig. 1e, Odds Ratio = 2.51, auPRC enrichment over baseline = 3.46). However, when we normalized nuclear size prior to running CAJAL, none of the ground truth genes were recovered, suggesting that CAJAL mixed size with shape when computing distances between nuclei.

In sum, TPCA representations are able to quantify different modes of shape variation (elongation, bean-shaped, triangular) amongst nuclei in the Schibler et al. screen. While well-known measures, such as eccentricity and roundness, captured half the variation in the data, additional data-driven features learned by TPCA were able to effectively recapitulate genes inferred by Schibler et al. Compared to CAJAL, TPCA was able to retrieve ground truth shape genes with higher confidence and without any confounding with nuclear size.

### KSTITCH recovers known shape–gene expression associations across multiple cell types in a Xenium dataset

Having established TPCA as a principled and effective approach to capture cellular or nuclear shape variability independent of cell size, we next develop a pipeline called KSTITCH (Fig 2a) to identify gene expression programs that co-vary with features of cellular shape in ISS-based (in situ sequencing) spatial transcriptomics datasets. Below we outline KSTITCH and test its ability to recover a known spatial gradient of nuclear eccentricity in keratinocytes as well as known relationships between morphology and gene expression in endothelial cells and monocytes/macrophages.

**Figure 2:**
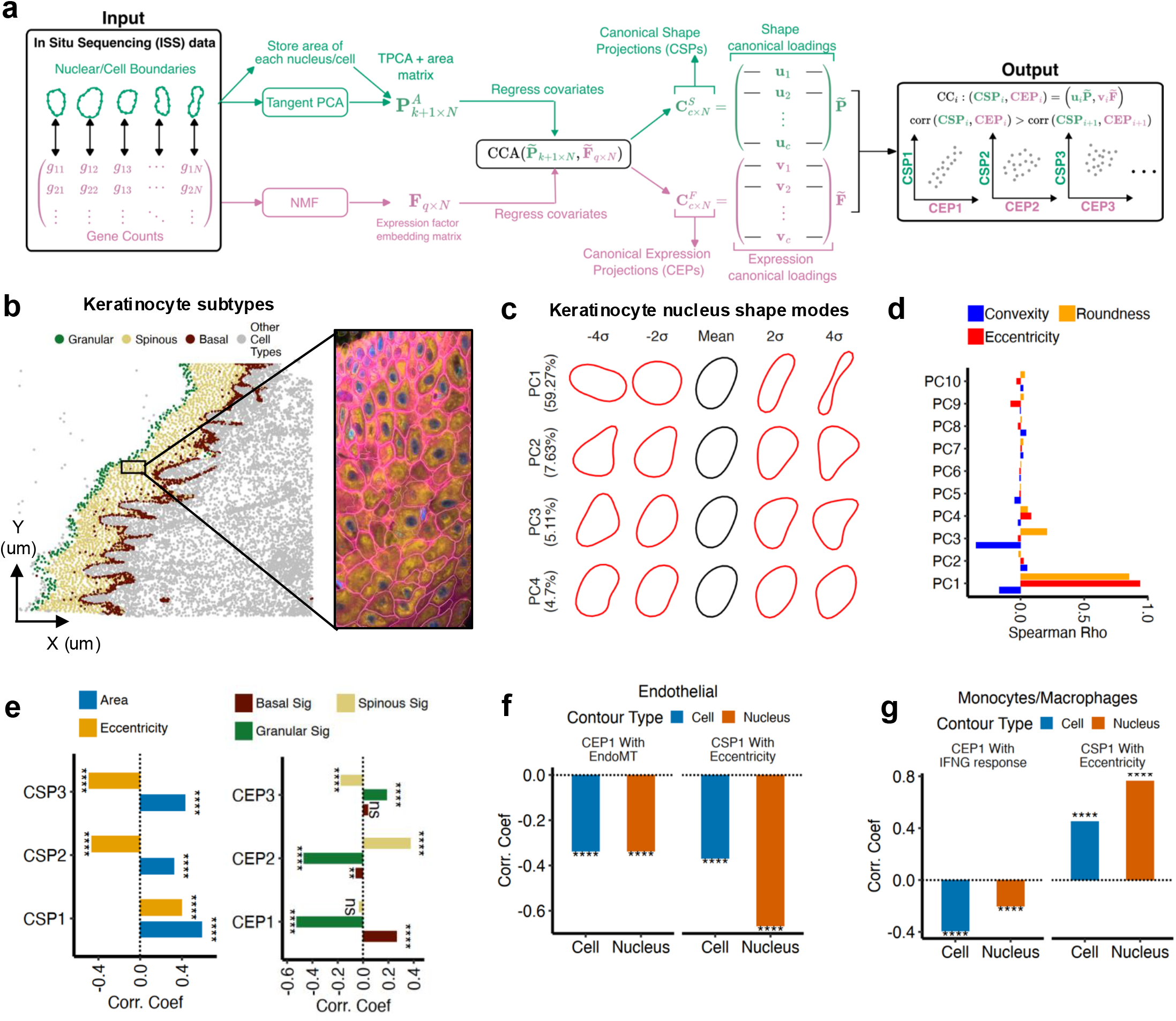
(a) STITCH pipeline to discover cell type-specific factors co-varying with cell or nuclear shape. First, shapes are represented in Tangent PCA space and the associated gene expression matrix is factorized using non - negative matrix factorization (NMF). Covariates are regressed from NMF and area-augmented PCA matrices (where each cell’s area is concatenated to its TPCA vector in the matrix) prior to computing CCA between them. This yields shape (u) and expression factor (v) loading vectors, which represent combinations of area and shape PCs with combinations of factors, respectively. These are used to compute canonical shape projections (CSPs) and canonical expression projections (CEPs), where the first canonical correlation (CC1) represented by (CSP1,CEP1) maximizes correlation between shape and gene expression. (b) Spatial plot of cells in Xenium Prime dermal melanoma dataset with keratinocyte subtypes labelled. Inset shows cell segmentation boundaries of keratinocytes (in purple), nuclear stains (blue) with associated boundaries (in red) and interior protein stain (orange). (c) Estimate of the mean shape of all keratinocyte nuclei (black), with shapes in red sampled between +4 and −4 standard deviations from the mean shape along each PC direction. Variance explained by each PC is shown in brackets. (d) Correlation of each shape PC derived from keratinocyte nuclei with the convexity, eccentricity and roundness (or circularity) of nuclei. (e) Left: Correlation of keratinocyte nuclear CSPs 1-3 with nuclear area and eccentricity. Right: Correlation of keratinocyte nuclear CEPs 1-3 with basal, spinous and granular keratinocyte signatures. (f) Left: Correlation of cellular (blue) and nuclear (red) CEP1 scores of endothelial cells with EndoMT signature. Right: Correlation of cellular and nuclear CSP1 of endothelial cells with cellular and nuclear eccentricity, respectively. (g) Left: Correlation of cellular (blue) and nuclear (red) CEP1 scores of monocytes/macrophages (or Mo/Macs) with Hallmark Interferon - γ response signature. Right: Correlation of cellular and nuclear CSP1 of Mo/Macs with cellular and nuclear eccentricity, respectively. **** p < 10^−4^, *** p < 10^−3^, ** p < 10^−2^, * p < 0.05.

The input to KSTITCH consists of paired data: gene expression count data and cellular (or nuclear) contour landmarks, for each cell derived from an ISS dataset. In the first step, cellular contours are embedded using TPCA while gene expression counts are embedded using the integrative non-negative matrix factorization (iNMF) algorithm from the LIGER package [24]. We concatenate the area of each cell or nucleus to its TPCA embedding vector to create an area-augmented TPCA embedding. In the second step, a canonical correlation analysis (CCA) is performed between expression-based NMF embeddings and area-augmented TPCA embeddings. We regress technical covariates (Methods) from the NMF and area-augmented TPCA embeddings depending on the ISS technology used. In CosMx datasets, we regress the total RNA count per cell and the distance of the cell from the edge of the field-of-view (FOV, when run on CosMx datasets), while the cell segmentation method employed and total RNA count per cell are regressed in Xenium datasets. CCA computes shape and expression canonical loadings—linear combinations of residualized shape PCs (and area) and expression factors, respectively—that maximize the correlation between canonical shape projections (CSPs) and canonical expression projections (CEPs) scores across cells. Each canonical correlation (CC) is represented by a CSP-CEP pair, with the first canonical correlation (CC1) representing the highest correlation between the respective CSP and CEP scores. Later canonical correlations represent CSP-CEP pairs that capture smaller correlations between shape and gene expression.

We analyzed a publicly available 10X Xenium Prime ISS dataset of a single FFPE (formalin-fixed paraffin-embedded) primary dermal melanoma sample. The dataset consisted of 112,551 cells whose expression was profiled using a panel of 5,006 genes. We processed this data using Seurat (v5.3.0, see Methods) to infer the identities of each cell based on their expression of cell type-specific marker genes (Supp. Fig. 2a,b). Certain cells were labelled “Unclassified” owing to low RNA counts (Supp. Fig. 2c) from that region. We further sub-clustered keratinocytes into granular, spinous and basal layers (Fig. 2b, Supp. Fig. 2d, Methods) based on the expression of known marker genes. We first computed TPCA embeddings of keratinocyte nuclear contours based on 50 sampled contour landmarks and retained the first 10 shape PCs capturing 90% of shape variation for downstream analyses. Fig. 2c illustrates the shape variation captured by the first four shape PCs. Similar to trends in the Schibler et al dataset, nuclear shape PC1 (capturing 59% of variation) varied with eccentricity and roundness (Fig. 2d). We then computed NMF embeddings (k = 10 factors) of keratinocytes using LIGER and ran KSTITCH to find canonical correlations of gene expression factors with nuclear area-augmented TPCA embeddings. We used a linear mixed model (see Methods) to remove technical variation associated with the cellular segmentation method used to determine cell boundaries and total RNA abundance from both shape PCs and expression factors prior to running KSTITCH. Though total RNA abundance variation within a spatial transcriptomics sample can represent tissue structure [25], the presence of low RNA counts from the “Unclassified” region in this dataset indicated that RNA count variation across cells in this dataset also arose from technical variation.

Spinous keratinocytes are known to have more rounded, i.e., less eccentric nuclei than granular and basal keratinocytes [26]. We therefore tested if the top canonical correlations represented correlations between nuclear eccentricity and transcriptional signatures of basal, granular, or spinous keratinocytes. The correlation coefficients associated with the CCs 1 to 3 from keratinocytes were 0.25, 0.16 and 0.1. The CC1 correlation coefficient was significant under both spatially restricted (p *<* 0.001, Methods) and global permutation nulls (p *<* 0.011), whereas later CCs were not formally tested for significance (Methods). As expected, CSP1 was positively correlated with nuclear eccentricity, while CEP1 was positively correlated with the basal signature and negatively correlated with the granular signature, indicating that CC1 captured a morphology–expression contrast between basal and granular keratinocytes (Fig. 2e). CSP2 was negatively correlated with eccentricity, whereas CEP2 was positively correlated with the spinous signature and negatively correlated with the granular signature, suggesting that CC2 distinguished spinous keratinocytes with more rounded nuclei from granular keratinocytes with more eccentric nuclei (Fig. 2e). Importantly, correlations between CEPs and keratinocyte signature scores, as well nuclear CSPs and eccentricity, remained statistically significant after accounting for total RNA abundance and cellular segmentation method, with only minor changes in effect size (Supp. Table 1A,B).

Next, we ran KSTITCH separately on nuclear and cellular contours of monocytes/macrophages (Mo/Macs) and endothelial cells to test whether the leading canonical correlations recovered known patterns of morphology–transcription covariation. As in the keratinocyte analysis, technical covariates were removed prior to running KSTITCH. Nuclear and cellular shape PC1 was associated with eccentricity in both cell types (Supp. Fig. 2e). Across nuclear and cellular analyses, CC1 correlation coefficients ranged from 0.18 to 0.4 and were significant under both spatially restricted and global permutation nulls (p *<* 0.05). Endothelial cells in tumors undergo endothelial-to-mesenchymal transition (EndoMT), a process accompanied by cellular and nuclear elongation [27]. Consistent with this relationship, endothelial nuclear CEP1 was negatively correlated with an EndoMT gene signature [28], while nuclear CSP1 was negatively correlated with eccentricity (Supp. Table 1C). The same pattern was observed for cellular elongation. Similarly, in Mo/Macs, CC1 recovered the known association between increased IFN*ω* signaling and decreased nuclear and cellular eccentricity (Supp. Table 1D) [2]. In both cell types, correlations between CEP1 and the corresponding gene-signature scores remained significant after accounting for total RNA abundance and cellular segmentation method, with only minor changes in effect size (Supp. Table 1C,D).

KSTITCH thus recovered the known spatial variation in nuclear eccentricity across keratinocyte layers, as well as known relationships between nuclear and cellular eccentricity and EndoMT or IFN*ω* signaling in endothelial cells and monocytes/macrophages, respectively. These associations persisted after accounting for total RNA abundance and segmentation-related variation.

### KSTITCH reproducibly recovers shape changes associated with fibroblast activation in a melanoma CosMx dataset

Our keratinocyte, endothelial, and monocyte/macrophage analyses involved shape and gene expression measurements in a single ISS slide. We then ran KSTITCH in a multi-sample cohort assayed using a different ISS platform (CosMx) to evaluate if (a) TPCA embeddings were robust to inter-slide variation, and (b) if KSTITCH could infer similar canonical correlations across samples. We first tested KSTITCH on fibroblasts as they are known to change shape and transcriptional state in response to tumor-derived signals [29].

We applied KSTITCH to a published ISS dataset generated using the Nanostring CosMx platform from ten melanoma patients (Love et al) with nevi (a benign melanocytic lesion), primary melanoma, or metastatic melanoma. These samples were imaged in three separate slides or batches, with each slide containing a primary, nevus, and metastatic sample. The CosMx platform profiled the expression of 968 genes in each cell and provided contour coordinates for each cell. Fig. 3a visualizes the spatial locations of pre-annotated malignant cells and microenvironmental cell types from a single sample (“Metastasis 1”) as well as the contours of each cell as determined by the CosMx segmentation pipeline in the inset. Unlike the Xenium dataset in Fig. 2, this dataset did not provide nuclear contours. Eight cell types were present across all samples (Supp. Fig. 3a), with malignant cells, fibroblasts, and macrophages/dendritic cells being the most abundant (Supp. Table 2A).

**Figure 3.**
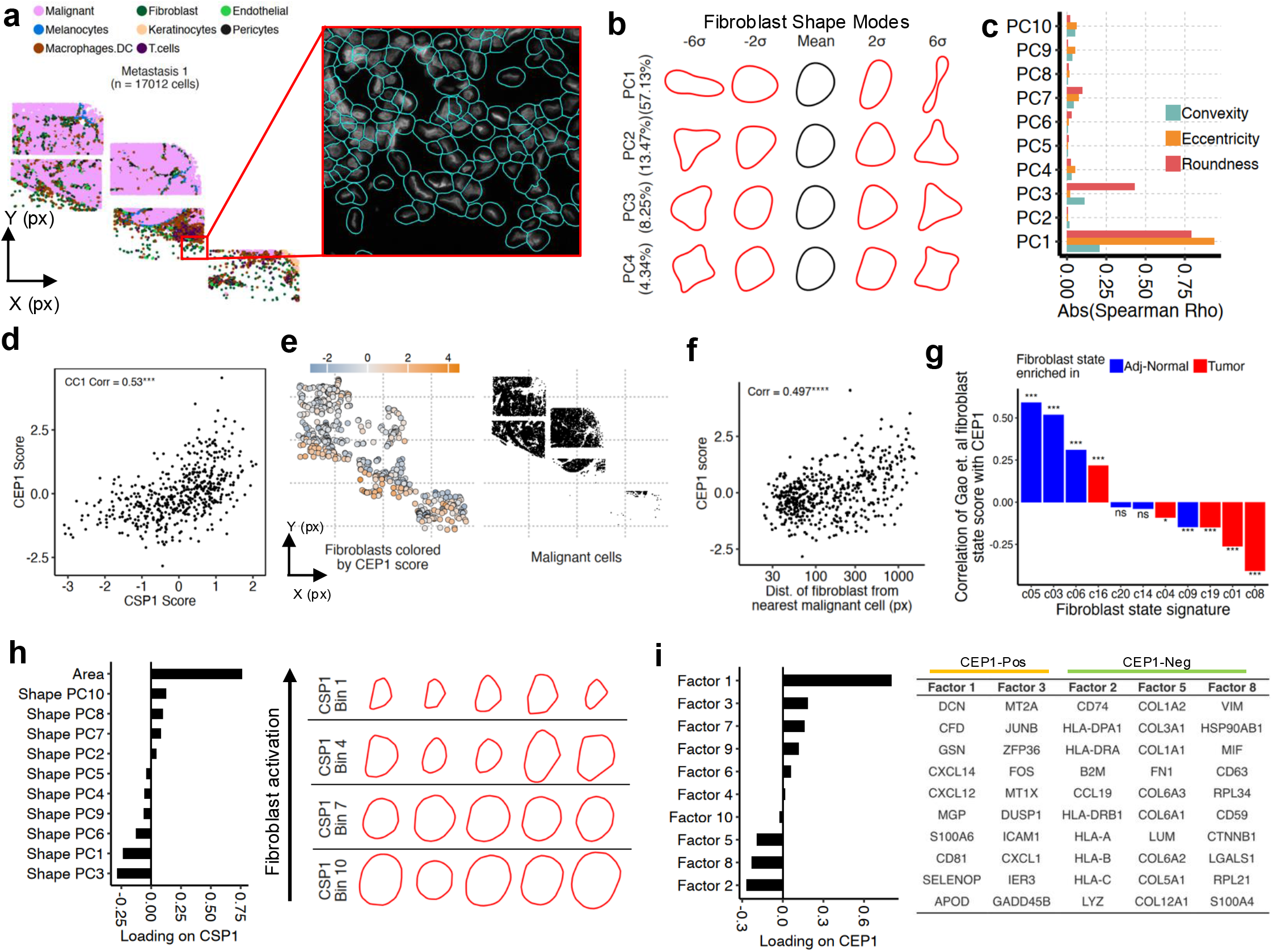
(a) Spatial coordinate plot of melanoma sample “Metastasis 1” from Love et. al melanoma cohort. Inset shows cell boundaries (in blue) of a selected region of the sample drawn by CosMx onboard segmentation method. (b) Estimate of the mean shape of all fibroblast cell boundaries (black), with shapes in red sampled between +6 and −6 standard deviations from the mean shape along each PC direction. Variance explained by each PC is shown in brackets. (c) Correlation of each shape PC derived from fibroblast cell boundaries with the convexity, eccentricity and roundness (or circularity) of nuclei.(d) Scatter plot of CEP1 and CSP1 scores of fibroblasts from sample “Metastasis 1”. **** p < 10^−4^ (e) Spatial locations of fibroblasts (left, colored by CEP1 score) and malignant cells (right) in the sample “Metastasis 1”. (f) Scatter plot of each fibroblast’s CEP1 score (x-axis) and its corresponding distance from the nearest malignant cell (y-axis) in the sample “Metastasis 1”. (g) Correlation of fibroblast CEP1 scores with expression of tumor-associated and adjacent-normal fibroblast signatures from Gao et. al. (h) Left: Loading of cell area and Shape PCs CSP1. Right: Five fibroblast cells from CosMx data at different values of CSP1 (bins along rows, with Bin 1 representing lowest CSP1 score). (i) Left: Loading of each factor on CEP1. Right: Top ten genes of each fibroblast gene expression factor, where high expression of factors 1 and 7 is associated with increased CEP1 while high expression of factors 2,5,8 are associated with decreased CEP1.

We ran TPCA and LIGER iNMF on fibroblasts across all samples and retained the top ten shape PCs of each cell type for further analyses. Both shape PC-based embeddings (Supp. Fig. 3b) and gene expression NMF embeddings of fibroblasts pooled across samples (Supp. Fig. 3c) did not contain any noticeable batch effects (average kBET [30] *p*-value *>* 0.9). Shape PC1 of fibroblasts captured 57% of variation and represented changes in cellular elongation (Fig. 3b,c) while PCs 2 and 3 represented other modes of shape variation.

Though TPCA and NMF embeddings were computed after pooling fibroblasts across samples, we first ran the CCA step in KSTITCH on fibroblasts from a single sample (“Metastasis 1”). Total RNA count per cell and the distance of each cell from the nearest field-of-view edge were regressed from NMF and TPCA embeddings prior to running KSTITCH to account for shape and total RNA count changes that can occur at the field-of-view edges in CosMx data [19]. Fig. 3d demonstrates the correlation between fibroblast CEP1 and CSP1 in this sample (R = 0.53, p *<* 0.001 under global and spatial permutation nulls). Interestingly, we found that fibroblast CEP1 scores were correlated with distance from the nearest malignant cell (Fig. 3e,f), which was in line with experiments linking tumor-derived signals to fibroblast shape and transcriptional changes [29]. Additionally, we correlated CEP1 scores with the expression of signatures of fibroblasts from Gao et al [31] found in tumors and normal tissue adjacent to tumors (“adjacent-normal”) and found CEP1 to be positively correlated with adjacent-normal fibroblast states and negatively correlated with tumor-associated states (Fig. 3g). This was consistent with the CEP1 scores of fibroblasts increasing in proportion to their distance from malignant cells.

We next inspected the loadings of cell area and individual shape PCs on CSP1 to determine which morphological features contributed to the expression-associated shape axis (Fig. 3h, left). High CSP1 scores were dominated by a strong positive contribution from cell area, whereas low CSP1 scores were associated with negative loadings on shape PCs 1 and 3. Consistent with the morphological variation represented by these PCs, fibroblasts with low CSP1 scores—which were located closer to malignant cells—were more eccentric and triangular, as confirmed by visual inspection of fibroblast contours across the CSP1 range (Fig. 3h, right)

We then inspected the loadings of gene expression factors on CEP1 (Fig. 3i, Supp. Table 2B) to identify transcriptional programs associated with these morphology changes. High expression of Factors 1 and 3 was associated with high CEP1, whereas high expression of Factors 2, 5 and 8 was associated with low CEP1. Factor 1 significantly overlapped (p *<* 0.001, permutation null) the c05-PI16 fibroblast progenitor signature from Gao et al., which is enriched in adjacent-normal samples (Supp. Table 2C). Factors 2 and 5 significantly overlapped the c20-CD74 antigen-presenting myofibroblast and c04-LRRC15 myofibroblast signatures, respectively. This was consistent with a recent study [15] linking spatial variation in fibroblast CD74 expression to nuclear shape variation in a bladder cancer Visium spatial transcriptomics dataset. Together, these results associated the elongated and triangular morphology of fibroblasts near malignant cells in the “Metastasis 1” sample with myofibroblast-associated and CD74+ transcriptional states.

Finally, we ran KSTITCH on fibroblasts in the remaining nine samples to check if similar shape-gene relationships were captured by CSP1 and CEP1. The CC1 correlation coefficient was significant in each sample (p *<* 0.001 under both global and spatial permutation nulls) and varied between samples (median = 0.35, IQR = 0.31 – 0.42, Supp. Fig. 3d), with the fibroblast CEP1 scores in each sample being positively correlated with distance from malignant cells (Supp. Fig. 3e). A decreasing fibroblast CEP1 score in each sample was correlated with increasing expression of tumor-associated fibroblast signatures from Gao et al. (Supp. Fig. 3f) where significance across samples was assessed using a random-effects meta-analysis (Methods). These correlations remained significant even after technical covariates (field-of-view distance, total RNA abundance) were regressed from Gao et. al signature scores (Supp. Table 2D). Loadings associated with CEP1 and CSP1 from each sample revealed that increased expression of Factors 2, 5 and 8 were consistently associated with a low CEP1 (Supp. Fig. 3g) while increased cell area, and decreased shape PCs 1 and 3 were associated with high CSP1 (Supp. Fig. 3h). Statistically, CEP1 and CSP1 canonical loading vectors were significantly aligned between samples (mean inter-sample cosine similarity of 0.79 and 0.52, respectively, *p <* 0.001 with a permutation null). Thus, KSTITCH reproducibly recovered a similar fibroblast morphology–expression relationship across all 10 independent patient samples in the Love et al. CosMx cohort.

### KSTITCH links mesenchymal-like melanoma cell states to increased cellular area

In contrast to fibroblasts, mutational differences between malignant cells across tumors increase transcriptional variation and potentially alter the link between transcriptome and morphology. This makes the robust inference of gene expression and shape relationships in malignant cells especially challenging. Having shown the ability of KSTITCH to reproducibly infer shape-expression relationships in fibroblasts, we assessed its efficacy on the more challenging problem of inferring shape-expression links in malignant cells. Melanoma cells are known to de-differentiate from a therapy-sensitive melanocytic (MEL) state towards a therapy-resistant mesenchymal (MES) state [32]. Since MES-like melanoma cells have been noted to have a larger surface area than MEL-like cells [33, 34], we checked if (a) KSTITCH can link MES gene expression to surface area across patients in the Love et al cohort, and (b) independently recover this link in a separate melanoma patient cohort [35].

We pooled malignant cells across the Love et al cohort and ran TPCA and iNMF. The first shape PC was associated with elongation and explained 50% of cell shape variation (Fig. 4a, Supp. Fig. 4a). We then ran KSTITCH separately on malignant cells from each patient. The CC1 correlation coefficient was significant in each sample (p *<* 0.001 under both global and spatial permutation nulls) and varied across samples (median = 0.25, IQR = 0.2–0.32, Supp. Fig. 4b). CSP1 and CEP1 loading vectors recovered from each sample were significantly aligned across samples with mean inter-sample cosine similarities of 0.9 and 0.37, respectively (*p <* 0.001), supporting the robustness of signals recovered by KSTITCH. Across samples, cell area was positively associated with CSP1 while shape PCs 1 and 3 were strongly negatively associated with CSP1 (Fig. 4b). Thus, an increase in CEP1 was associated with an increase in malignant cell area.

**Figure 4.**
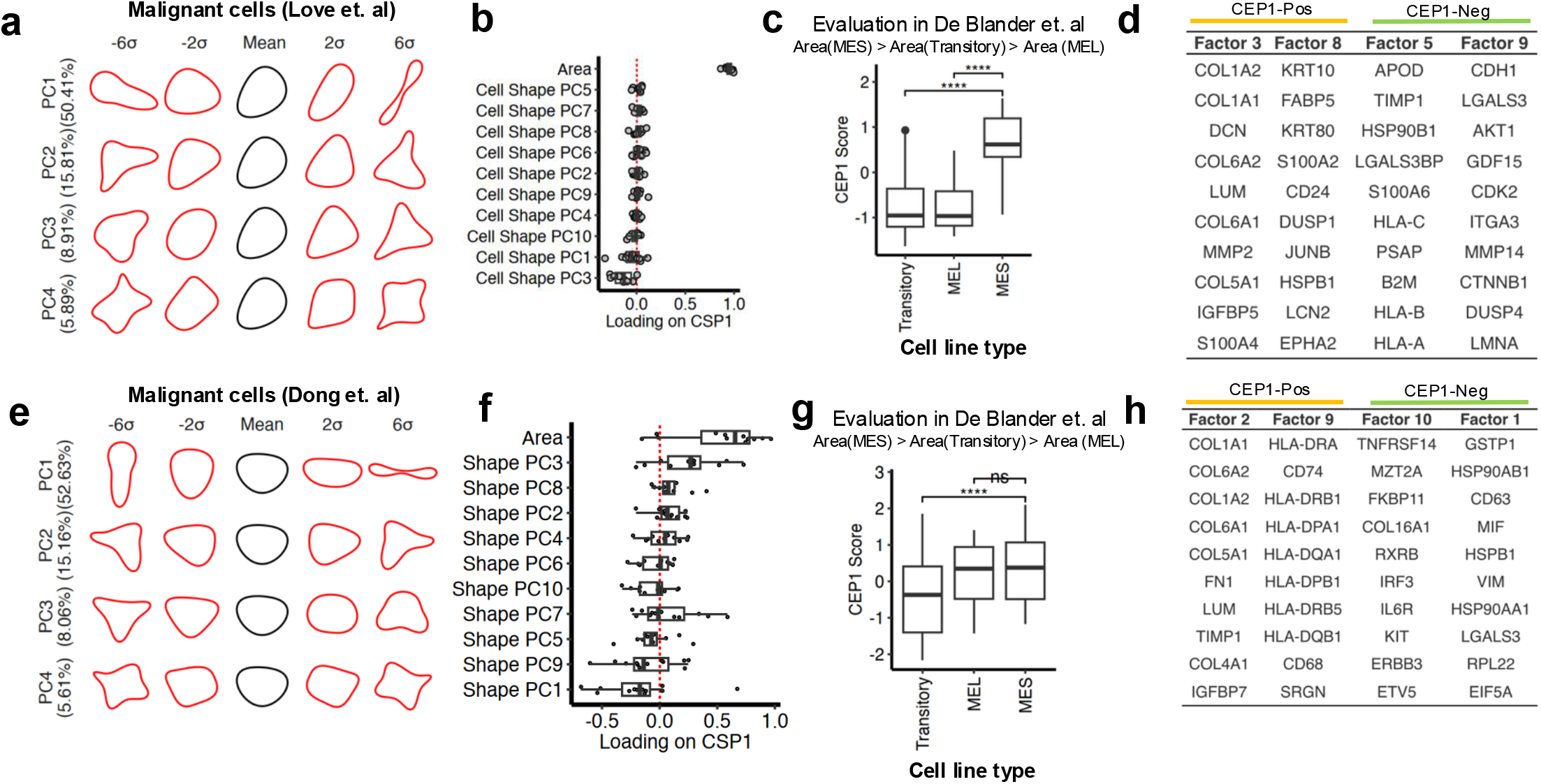
(a) Estimate of the mean shape of all malignant cell boundaries (black) from Love et. al cohort, with shapes in red sampled between +6 and −6 standard deviations from the mean shape along each PC direction. Variance explained by each PC is shown in brackets. (b) Loadings of each malignant cell shape PC and area with CSP1. Each dot is a different sample from Love et. al cohort. (c) Estimated CEP1 scores from each Love et. al sample in bulk RNA-seq of melanoma cell lines in MES, MEL and Transitory states. Mean 2D area of cell lines in each of these states was measured in De Blander et. al with MES cell lines having the highest area followed by Transitory and MEL cell lines. (d) Top ten genes of each malignant gene expression factor in Love et. al cohort, where high expression of factors 3 and 8 is associated with increased CEP1 while high expression of factors 5 and 9 are associated with decreased CEP1. (e) Estimate of the mean shape of all malignant cell boundaries (black) from Dong et. al cohort, with shapes in red sampled between +6 and −6 standard deviations from the mean shape along each PC direction. Variance explained by each PC is shown in brackets. (f) Loadings of each malignant cell shape PC and area with CSP1. Each dot is a different sample from Dong et. al cohort. (g) Estimated CEP1 scores from each Dong et. al sample in in bulk RNA-seq of melanoma cell lines in MES, MEL and Transitory states. Mean 2D area of cell lines in each of these states was measured in de Blander et. al with MES cell lines having the highest area followed by Transitory and MEL cell lines. (h) Top ten genes of each malignant gene expression factor in Dong et. al cohort, where high expression of factors 2 and 9 is associated with increased CEP1 while high expression of factors 1 and 10 are associated with decreased CEP1.

To test this association in an external dataset, we used each sample’s CEP1 canonical loading vector to compute CEP1 scores in a melanoma cell line panel (Methods) whose mean cell areas were measured by De Blander et al [34]. MES cell lines were found to have higher cell areas than MEL and Transitory cell lines in this study. As expected, we found that the MES cell lines had a higher CEP1 score than both the MEL and Transitory cell lines (Fig. 4c). We further checked if the highest-weight genes in factors associated with CEP1 overlapped known MES and MEL signatures. We picked the two expression factors with the highest loadings in CEP1 and computed the overlap of the 50 highest weighted genes in each factor (Fig. 4d, Supp. Table 3A) with melanoma de-differentiation signatures from Tsoi et al [32] and Andrews et al [36]. Factors 3 and 8, associated with high CEP1, significantly overlapped the Andrews et al MES signature and Tsoi et al undifferentiated-neural crest-like signature, respectively (p *<* 0.01, Supp. Table 3B, permutation null). Thus, genes ultimately underlying CEP1 represented known transcriptional differences between MES and MEL cell lines.

Finally, we tested if KSTITCH could recover the link between MES-like expression and increased cell area in a different (Dong et al, [35]) melanoma cohort. This cohort posed a greater challenge for our method, as it encompassed a broader range of disease stages (22 cutaneous melanoma samples, stages I-IV), treatment status (pre- and post-immunotherapy), metastatic sites, and fewer malignant cells per sample (median = 1,934 cells compared to median = 11,403 cells in Love et. al). Further, each patient sample was imaged in only a single field-of-view, which meant that it could not be regressed as a covariate from shape PCs and gene expression factors. We ran TPCA and NMF on malignant cells from this cohort independently of the Love et al cohort above. The recovered shape PC modes (Fig. 4e) closely resembled those recovered in the Love et al cohort, with PC1 covarying with cell eccentricity (Supp. Fig. 4d). We regressed total RNA abundance and the distance of each cell from the field-of-view margin from shape PCs and gene expression factors and ran KSTITCH separately on each sample. The range of CC1 correlation coefficients was lower than that seen in the Love et al cohort (median = 0.17, IQR = 0.15 – 0.21, Supp. Fig. 4b), with 11/22 samples having a significant CC1 correlation coefficient (*p <* 0.05) under both global and spatial permutation nulls. Cell area was positively associated with CSP1 (Fig. 4f) across samples, with shape PC1 being negatively associated with CSP1. The inter-sample alignment of CSP1 and CEP1 loading vectors across samples was lower than in Love et. al (mean inter-sample cosine similarities of 0.436 and 0.239, respectively) but was statistically significant (*p <* 0.001, permutation null). Nonetheless, despite this heterogeneity, CEP1 loading vectors from each sample successfully ranked MES cell lines above Transitory cell lines in the De Blander et al. panel (Fig. 4g,h), supporting cross-cohort reproducibility of the MES–area link.

We inspected the top factors associated with CEP1 in Dong et al, and found that Factor 2 (associated with high CEP1) significantly overlapped the Andrews et al MES signature while Factor 10 (associated with low CEP1) overlapped the Andrews et al MEL signature (Supp. Table 3C,D). More broadly, each factor recovered in the Dong et al cohort could be mapped to a factor recovered in the Love et al cohort (Supp. Table 3E).

These analyses demonstrated that KSTITCH reproducibly recovered a link between MES-like gene expression in malignant cells and cell area when run independently in two separate cohorts. Crucially, the canonical expression loadings recovered from both cohorts could be successfully mapped to a different transcriptomic protocol (bulk RNA-seq) where CEP1 scores co-varied with an independent measurement of cell area.

## Discussion

KSTITCH integrates cellular or nuclear morphology with transcriptomic data to discover interpretable patterns of covariation between morphology and gene expression. Notably, it relies on TPCA, a principled and well-established method of shape manifold analysis, to separate size (or area) from an object’s shape. We demonstrated the value of separating size from morphology by showing that TPCA was able to identify experimentally ascertained nuclear morphology-related genes with high specificity, while a previously published metric geometry-based approach – CAJAL, was driven largely by size rather than shape characteristics.

We also show that KSTITCH recovers known relationships between morphology and gene expression across keratinocytes, endothelial cells, and macrophages in a Xenium ISS dataset. In a melanoma patient cohort profiled using the CosMx ISS platform, KSTITCH inferred concomitant shape and transcriptional changes in fibroblasts proximal to malignant cells. The transcription-coupled morphology change discovered in fibroblasts involved changes in area but also changes in a shape feature (shape PC3) that was associated with a triangular shape on visual inspection that did not correlate with standard shape descriptors such as eccentricity and convexity. This relationship was recovered independently in fibroblasts from each patient sample, suggesting the reproducibility of KSTITCH and its ability to robustly detect shape features beyond those that are visually discernible in current practice. Our finding recapitulates recent observations of activated fibroblasts isolated from breast and pancreatic cancers having a triangular morphology [37, 38]. Finally, KSTITCH independently recovered an experimentally observed link between melanoma cell differentiation state and area [33, 34] across multiple patient samples as well as across cohorts.

The inferences made by KSTITCH are independent of several sources of technical variation such as (a) the dependence of cell morphology and total read count on distance from the field-of-view in CosMx datasets, and (b) differences in the segmentation method used to draw cell contours in Xenium Prime data. There are other factors that may still impact KSTITCH’s inferences. Transcript mis-assignment between neighboring cells due to imperfectly drawn cell contours [39, 40] could either induce spurious co-variation or lower true co-variation between gene expression and morphology. We minimize this confounding by restricting our analyses to those genes known to be robustly expressed in corresponding cell types in the Human Protein Atlas. This, however, deals with misassignments between neighbor cell pairs that are of different types but not those of the same type or in subtly different transcriptional states.

KSTITCH could be modified and extended in the future. TPCA relies on Kendall shape manifolds. An alternate approach to a Kendall shape manifold is one constructed via elastic shape analysis [41], where TPCA representations can capture more complex cell morphologies, such as seen in vitro, by also optimally repositioning landmarks along contours. Though the morphology–expression relationships recovered in our work appear to be approximately linear, any switch-like or threshold relationships, if present, would require nonlinear extensions such as DeepCCA or Kernel CCA [42] that can potentially recover such effects but do so at the cost of model interpretability. An extension to multi-modal shape and gene expression embedding could involve a MOFA-type approach [43] to construct a common latent space underlying both gene expression and TPCA representations of each cell. Ultimately, our work highlights the value of manifold-based shape representations in statistical analysis of cell morphology, in a manner that (1) disambiguates shape features from cellular or nuclear size explicitly, (2) is interpretable, and (3) is robust across ISS technologies.

## Acknowledgments

We thank Kerrie Marie, Victoria Sanz-Moreno, Etan Aber, Eldad Shulman and Eytan Ruppin for discussions, Baktiar Karim for analysis of the pathology slide associated with the Xenium Prime melanoma dataset. We thank Gianluca Pegoraro for providing data and discussions on the Schibler et al shape screen. We used OpenAI’s ChatGPT (v5.4) to assist with editing text in the abstract and methods sections of the manuscript.

This research was supported [in part] by the Intramural Research Program of the National Institutes of Health (NIH). The contributions of the NIH author(s) were made as part of their official duties as NIH federal employees, are in compliance with agency policy requirements, and are considered Works of the United States Government. However, the findings and conclusions presented in this paper are those of the author(s) and do not necessarily reflect the views of the NIH or the U.S. Department of Health and Human Services. This work utilized the computational resources of the NIH HPC Biowulf cluster (http://hpc.nih.gov). This work was supported by U.S. National Cancer Institute grants (ZIABC012176-01) to SH. SK is supported by National Institute of Neurological Disorders and Stroke Grant R01NS134842. YS is supported by Harvard Ludwig Center and the Mark Foundation for Cancer Research. TV is supported by a Research Scholar Grant, PF-24-1316850-01-CD, from the American Cancer Society, Harvard Ludwig Center and the Mark Foundation for Cancer Research. AS is supported by the National Science Foundation (NSF grant 2413748) and US Air Force Office of Scientific Research (AFOSR FA9550-23-1-0673).

## Methods

### Kendall Shape Space and Tangent Principal Component Analysis

Segmentation algorithms extracts contours of cell and nuclei from imaging data and represent each by a sequence of coordinates – called *landmarks* – along these closed contours. Our goal is to compare and interpret nuclear and cellular shape variation using statistical techniques that are invariant to rotations, translations, reflections, the ordering of landmarks, and differences in size. Since these differences can arise as artifacts of the imaging process, we treat them as *nuisance transformations* rather than meaningful biological variation. We note that nuclear and cellular size differences are a nuisance variable when comparing shapes but their *co-variation* with the transcriptome can be biologically meaningful [44] and are included separately in KSTITCH. Removal of these nuisance transformations embeds cells and nuclei in a shape space *S* that is a nonlinear manifold where each point on *S* represents a shape. We will first describe the landmark-based or Kendall’s shape analysis to form *S*, and then describe tangent principal component analysis (TPCA), an adaptation of the classical (Euclidean) PCA to the non-linear geometry of *S*. The term contour used below encompasses both nuclear and cellular contours. Implementation details for computing TPCA coordinates from contours is shown in the appendix (Algorithms 1—4).

#### Step 1: Embedding contours in *S*

We first lay out the steps used to remove nuisance transformations and to compute distances between them as elements of *S*. Let *X* = [*X*_1_ *X*_2_, …, *X*_*L*_] be a 2 × *L* matrix denoting *L* planar points along a contour. The effect of translation i.e., the location of a contour in an image, is removed by subtracting the mean from the landmarks, *i*.*e*., 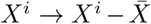, where 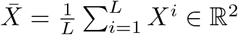. We impose invariance to cellular or nuclear size by rescaling the matrix *X*^*p*^ = *X/* ∥ *X* ∥ _F_, where ∥ *X* ∥ _F_ is the Frobenius norm of *X*. The resulting shape *X*^*p*^ (where superscript *p* denot resides in a pre-shape space which is a unit sphere of dimension 2*L* − 3. The remaining nuisance transformations to be removed are rigid rotation, reflection, and cyclic landmark order. These are removed in a pairwise manner using a given shape as a reference. Let *X*^*p*^,*Y* ^*p*^ ∈ *C* denote two pre-shapes i.e., centered and size-normalized landmark configurations for two contours, and let *C*_*m*_ denote the cyclic-shift operator that cyclically shifts the *L* landmarks by *m* positions, *i*.*e*., *C*_*m*_([*X*_1_, *X*_2_, …, *X*_*L*_]) = [*X*_*L−m*+1_, …, *X*_*L*_, *X*_1_, …, *X*_*L−m*_]. Let O(2) denote the set of all 2 × 2 orthogonal matrices. Then, the space resulting from the removal of cyclic shifts and rotation is denoted by *S* = *C /*(O(2) × *C*_*m*_) and is termed the shape space. Elements of are sets of type [*X*^*p*^] which contain all possible rotations, reflections, and cyclic shifts of *X*^*p*^ ∈ *C*. To rotate (or align) *Y* ^*p*^ to *X*^*p*^, we search over all cyclic shifts of the landmarks and, for each shift, compute the optimal rotation/reflection matrix by Procrustes alignment

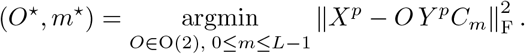

For each fixed shift *m*, the optimization over O(2) is computed as *O*^★^ = *V U* ^*T*^, where (*Y* ^*p*^*C*_*m*_)(*X*^*p*^)^*T*^ = *U* Σ*V* ^*T*^ is the singular value decomposition and *T* denotes the transpose. Then *Y* ^*p*★^ = *O*^★^*Y* ^*p*^*C*_*m*★_ is the optimally rotated and landmark-shifted shape with *X*^*p*^ as a reference. The geodesic distance between original contours *X* and *Y* is then *d*_s_(*X, Y*) ≜ cos ^−1^(*X*^*p*^,*Y* ^*p★*^).

#### Step 2: Tangent Principal Component Analysis

The first step in TPCA computation is defining the mean shape (or Fréchet mean) of a given set of shapes *{X*^(*j*)^,*j* = 1, …, *N}* ∈ *S* from a dataset after implementing the operations described above. Let *d*_s_(*X*^(*i*)^,*X*^(*j*)^) be the shape distance between arbitrary contours *X*^(*i*)^ and *X*^(*j*)^. Then, we define the mean shape of this sample set as:

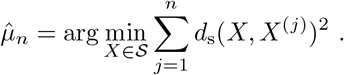

There is a well-known iterative algorithm to find this minimum and is presented in the appendix (Algorithm 3). Since the shape space *S* is nonlinear, one cannot use the Euclidean residuals 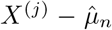 to compute their covariances. Instead, one uses a mapping called logarithmic map that maps the observed shapes *{X*^(*j*)^*}* to the flat tangent space of 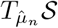 at 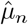 (the formula is provided in the appendix in Algorithm 3). The resulting tangent vectors 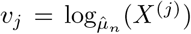 are elements of 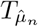 and can be used to define sample covariance using 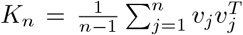, where the superscript *T* denotes the transpose. Using singular value decomposition (SVD), one can decompose *K*_*n*_ = *U* Σ *U* ^*T*^, and obtain a natural tool for dimension reduction and shape representation. Let *Ũ* denote the first *k* columns of the orthogonal matrix *U*, then 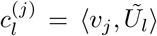 is the *l*^*th*^ principal coefficient or shape PC score of the shape *X*^(*j*)^. Each shape *X*^(*j*)^ is conveniently represented by a vector 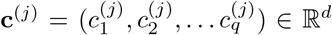 with the diagonal entries of Σ providing the variance explained by each shape PC. Together, these steps — estimating the mean shape 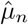, mapping each observed shape to a tangent vector *v*_*j*_, estimating the tangent-space covariance *K*_*n*_, and decomposing this covariance matrix to obtain *{***c**^(**1**)^, **c**^(**2**)^, …, **c**^(**N**)^ *}* —constitute tangent principal component analysis (TPCA).

### TPCA and CAJAL analysis of CRL1474 fibroblast nuclear images from Schibler et al

In Schibler et al, each well in a 384-well plate consisted of CRL1474 fibroblasts where the expression of a single protein was inhibited by one of three small interfering RNAs (siRNAs) targeted to different parts of its sequence.

Nuclei were imaged in multiple fields of view from each well in three channels – DAPI expression, *LMNB* expression and *LMNA* expression. Each siRNA perturbation was carried out in two biological replicates. In total, 867 nuclear proteins were perturbed with each siRNA perturbation being carried out in two biological replicates.

We used images in the DAPI channel to segment nuclei using cellpose v3.0.5 [45] where DAPI intensities were scaled to range [0, 255] in each image prior to running cellpose. The nuclear diameter in each image was estimated by cellpose by setting the `--diameter` option to `0` with the pre-trained nuclear model being employed for segmenting nuclei. Nuclei whose segmentation masks intersected the borders of the image were removed and contours of remaining nuclei were extracted using the Python OpenCV library (v4.12.0). Since inspection of segmentation masks and raw images revealed that some masks represented nuclei from two different cells that overlapped each other in the z-plane while others potentially represented micro-nuclei, we computed the area of every segmentation mask and only retained nuclei whose segmentation masks lay between the 5^*th*^ and 95^*th*^ percentile of mask areas within a given image. A total of *N* = 2, 011, 829 nuclei were retained for further analyses with 2059 to 2614 nuclear contours (inter-quartile range) obtained per perturbation (inclusive of non-targetting siRNA probes that represented nuclear contours from unperturbed cells).

For each nuclear contour, we used linear spline interpolation to obtain 50 landmarks prior to running TPCA on all nuclei and retained the first 10 shape PCs for further analysis. In order to detect genes whose perturbation alter nuclear shape, we first constructed a reference distribution of nuclear shape variation amongst unperturbed nuclei by computing the distance of PC vectors of all unperturbed nuclei from their medoid. We then computed all the distances of nuclei from each perturbation to the medoid of unperturbed medoid and compared it with the reference distribution using a one-sided Wilcoxon rank-sum test with the alternative hypothesis that distances of perturbed nuclei from the unperturbed medoid was larger than that of the reference distribution. We then employed a criterion similar to the one followed by Schibler et al to detect shape-related genes, where a gene was considered to perturb nuclear shape if the distances of perturbed nuclei from the unperturbed medoid was significantly greater (*FDR <* 0.1) than that of unperturbed nuclei for at least two of the siRNAs that targeted the gene. We repeated this calculation after excluding the first shape PC or the first two shape PCs to determine whether these features signifcantly reduced the overlap of shape genes inferred by our approach with those inferred by Schibler et al.

*CAJAL*: As per instructions from the CAJAL documentation, it was run using the contour landmarks obtained from cellpose without any interpolation. CAJAL was first run on nuclei from the unperturbed (or negative control) condition to determine the unperturbed medoid. CAJAL was then run to compute the exact Gromov-Wassterstein distance between each perturbed nucleus and the unperturbed medoid. In order to run CAJAL with size-normalized nuclei, we divided each contour landmark coordinate by the Frobenius norm of the landmark matrix representing each nucleus and ran the CAJAL pipeline as outlined above.

### Description of KSTITCH pipeline

In addition to cellular or nuclear contours associated with *N* cells in a slide, spatial ISS data yields an expression matrix **E**_|*G*|×*N*_ that contains transcript counts of genes *G* = *{g*_1_, *g*_2_, … *}* for each cell. We utilize non-negative matrix factorization (NMF) to reach Euclidean representations of gene expressions and analyze them jointly with the shape TPCA representations.

The overall pipeline consists of three steps. In Step 1 of the pipeline, for each cell, we applied NMF on the gene expression matrix to obtain a factor representation and applied TPCA on nuclear or cellular contours to compute TPCA representations. The cell area (a scalar quantity) is concatenated to its TPCA representation vector to generate an area-augmented TPCA vector. In Step 2, we regress out any technical covariates from the joint factor and TPCA representations. In Step 3, CCA is performed between the regressed factor and area-augmented TPCA representations to obtain canonical shape and expression loading matrices that are then used to compute canonical shape projections (CSPs) and canonical expression projections (CEPs) of cells. If cells of different types are present in the dataset, steps 1-3 are run separately on each cell type.

#### Step 1: NMF and TPCA

If the *c*^*th*^ cell type consists of *m* cells, we decompose 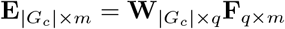 using NMF of rank *q*, where **F**_*q×m*_ are the factor embeddings of each cell and 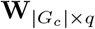 are the weights of each gene in each factor. We use the LIGER integrative NMF (iNMF) implementation [24] for this computation and set *q* = 10 factors as a default. We confirmed the reproducibility of the factors recovered as outlined by Kotliar et al [46]. Factors are interpreted by inspecting 50 genes with the highest loadings in **W**. *G*_*c*_ is the gene subset specific to that cell type, *i*.*e*., genes that are expressed in at least 1% of cells and have an expression level of at least 10 nTPM in the corresponding cell type in the human protein atlas [47]. This filtering step is done to remove genes whose transcripts are incorrectly assigned from neighboring cells due to cell segmentation error [40, 39]. In addition to NMF, we apply TPCA on the same cells to obtain a TPCA embedding matrix **P**_*k × m*_ = [**c**^1^ **c**^2^ … **c**^*N*^] as described before, where *k* represents the top *k* shape PCs that capture ~ 90% of variance. The area of each cell, denoted by *A ϵ* R_+_, is computed and concatenated to each TPCA vector to obtain an area-augmented TPCA matrix 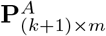.

**Step 2: Covariate regression:** We regress two sets of covariates from **P**^*A*^ and **F**, irrespective of the ISS platform used to generate the data, with platform-specific covariates added as necessary. The basic covariates considered are the total RNA count per cell, 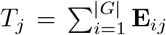, computed across all genes and not just cell type-specific genes. *T*_*j*_ is regressed from each expression factor, and each shape PC. Formally, the following linear model is implemented in R using the `glm` function for covariate regression –

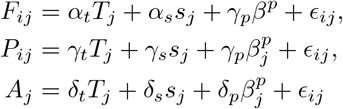

where ϵ_*ij*_ *~N* (0,*σ* ^2^), *F*_*ij*_ and *P*_*ij*_ represents the coordinate or score of expression factor *i ϵ{*1, …, *q}* and shape PC *i ϵ{*1, …, *k}* in cell *j. A*_*j*_ is the area of cell or nucleus *j*. 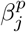 refers to platform-specific covariates that are added based on the ISS platform used. *s*_*j*_ is an optional categorical variable that represents the slide or sample index in which the cell is profiled if a multi-sample cohort is analyzed. In CosMx data, 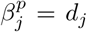 where *d*_*j*_ is the distance of the *j*^*th*^ cell from the field-of-view edge. We also add the field-of-view ID as a random e!ect to the regressions when a given sample is profiled across multiple fields of view. In Xenium datasets where cell segmentation staining is available, β^*p*^ is a categorical variable whose value signifies if the cell contour was drawn based on either nuclear expansion, interior protein staining or a cell membrane stain.

**Step 3: CCA:** CCA is performed between the residualized matrices 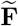 and 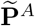 obtained from Step 2, where 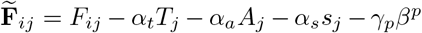 and 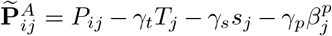. CCA learns canonical loading matrices **U** and **V**, where the rows of **U** are area-augmented TPCA loading vectors and the rows of **V** are expression loading vectors, such that the paired projections 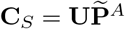 and 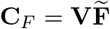 are maximally correlated:

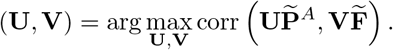

The rows of **U** and **V** are the canonical shape and expression loading vectors, respectively, and the rows of **C**_*S*_ and **C**_*F*_ are the canonical shape projections (CSPs) and canonical expression projections (CEPs), respectively.

#### Permutation testing of canonical correlations

Significance of the first canonical correlation (CC1) was assessed using both global and spatially restricted permutation null models. For the global null, cell identities associated with the expression-factor matrix were randomly permuted prior to CCA, thereby disrupting cell-specific correspondence between expression and morphology. For the spatially restricted null, cells were first partitioned into spatial bins based on their centroid coordinates, and expression-factor vectors were permuted only among cells within the same spatial bin to preserve coarse spatial structure while disrupting cell-specific morphology–expression pairing. The purpose of this test is to control for the baseline expectation that spatially proximal cells will have similar expression patterns as well as morphologies without any direct link between the two. Empirical *p*-values were calculated as the fraction of permuted CC1 values greater than or equal to the observed CC1. Permutation testing was restricted to CC1 because naive permutation inference for subsequent canonical correlations can yield inflated error rates unless canonical modes are tested sequentially after accounting for variance explained by preceding modes [48]. Therefore, later canonical correlations were not formally tested for significance.

### Processing of Xenium Prime primary dermal melanoma dataset

Processed data consisting of the gene expression matrix of all cells and their associated cellular and nuclear contours were downloaded from the 10x website https://www.10xgenomics.com/datasets/xenium-prime-ffpe-human-skin. We removed cells from analysis that had either had zero or more than one nuclear contours associated with it.

In order to label each cell type, we used the `SCTransform` function in `Seurat` (v5.3.0) to normalize read counts across all cells. We then calculated a PCA embedding of all cells based on the normalized and scaled expression of the 2000 most variable genes, and clustered cells using the`FindClusters` function of `Seurat` with default parameters based on the first 30 PCs. We then relied on the expression of marker genes shown in Supp. Fig. 2b to assign labels to each cluster. An expert pathologist examined the associated H&E image to determine areas of poor RNA preservation (where most were labelled as “Unclassified”) and verified gross cell type assignments of immune infiltrates, malignant cells and keratinocytes within the image.

### Adjustment of gene-signature scores for technical covariates

To assess whether associations between canonical expression projections (CEPs) and published gene-signature scores in keratinocytes, endothelial cells and monocytes/macrophages were robust to technical variation, signature scores were residualized against total RNA abundance and cellular segmentation method using linear regression. Correlations between CEP scores and the residualized signature scores were then recomputed and compared with the corresponding unadjusted correlations. Associations were considered robust when their direction was preserved and e!ect sizes changed only minimally after covariate adjustment.

### Processing of Love et al and Dong et al Nanostring CosMx dataset

We downloaded expression counts and cellular contours for each sample in the Love et al cohort from the Dryad repository https://datadryad.org/dataset/doi:10.5061/dryad.ksn02v7b1, and processed the data using Seurat. We ran SpaceTrooper [19] to remove the lowest quality cells at the edge of each field-of-view. Cells were annotated using the `cluster` column in the annotation file, with all sub-clusters of each cell type grouped into fibroblasts, macrophages/DCs, pericytes, endothelial cells, malignant cells, T cells, keratinocytes and melanocytes. For each cell type, we grouped cells across samples and ran TPCA and retained the top 10 shape PCs for further analysis (*n* = 50 landmarks were interpolated for each cellular contour). We ran NMF on cells of each type separately using *q* = 10 factors. KSTITCH was run with total RNA count per cell, cell area, and CosMx-specific covariates as described above.

We downloaded expression counts and cellular contours for each sample in the Dong et al cohort from the Zenodo repository https://zenodo.org/records/14708000, and processed the data using Seurat and used SpaceTrooper to remove low-quality cells. Since the average number of non-malignant cells in each sample was too low (*<* 100 per sample), only malignant cells were retained for further analyses. NMF and TPCA were run using the same parameters as for Love et al. KSTITCH was run with total RNA count per cell, cell area, and CosMx-specific covariates as described above.

### KSTITCH analysis of fibroblasts from Love et al cohort

The presence of batch e!ects amongst fibroblasts was tested using kBET [30]. KSTITCH was run separately on area-augmented TPCA and NMF matrices of fibroblasts subsetted to contain cells from each sample in the Love et al. cohort. The distance of each fibroblast from the nearest cell of every other cell type was computed using the nn2 function in the annoy package in R. Signatures of fibroblast states were obtained from Gao et al. [31] and subset to contain genes profiled in the 968-gene CosMx panel. Of the 20 fibroblast signatures derived by Gao et al. from fibroblasts across multiple cancer types, including melanoma, we analyzed those that were (a) enriched in normal tissues adjacent to a tumor (“adjacent-normal”) or in tumor samples and (b) enriched in skin or present across multiple organ sites including skin.

CEP1–signature correlations were computed separately within each Love et al. sample. To obtain a single cross-sample e!ect-size and significance estimate for each Gao et al. fibroblast signature, sample-specific correlation coe”cients were transformed using Fisher’s *r*-to-*z* transformation and combined using a random-e!ects meta-analysis with sampling variance 1*/*(*n* 3), where *n* is the number of cells in the corresponding sample. The pooled estimates were transformed back to correlation coe”cients for reporting, and meta-analysis *p*-values were adjusted across signatures using the Benjamini–Hochberg procedure. To assess robustness to technical covariates, Gao et al. signature scores and CEP1 scores were separately regressed for total RNA abundance, distance from the nearest field-of-view margin, and field-of-view identity, after which the sample-specific correlations and random-e!ects meta-analysis were recomputed.

To assess the spatial association between fibroblast CEP1 and surrounding cell types, we computed, for each fibroblast, the Euclidean distance to the nearest cell of each background cell type. Within each sample, Pearson correlations were then computed between fibroblast CEP1 scores and distance to each background cell type. To summarize these associations across samples, sample-specific correlation coe”cients were transformed using Fisher’s

*r*-to-*z* transformation and meta-analytic estimates were derived as described above.

### KSTITCH analysis of malignant cells in Love et al and Dong et al cohorts

KSTITCH was run independently on malignant cells of each sample in Love et al and Dong et al cohorts based on area-augmented TPCA and NMF matrices. *q* = 10 factors were computed from each cohort, with **W**_*L*_ and **W**_*D*_ representing NMF gene weight matrices from each cohort.

We downloaded RNA-seq read counts from the GEO database (GSE134432) for all cell lines except NCL (neural crest-like) cell lines, which were not publicly available. We obtained a normalized read count matrix for each cell line by scaling read counts to counts-per-million (CPM), after which each gene’s expression variance was unit-scaled. We used the annotation of cell lines as MES, MEL and Transitory provided by De Blander et al.

In order to compute canonical expression projections from bulk RNA-seq data of cell lines profiled in De Blander et al, we need to first compute an approximate factor embedding of each cell line after which the factor embeddings are multiplied by the canonical expression loading matrix. Let **X** be the normalized read count matrix obtained from cell lines where the columns represent cell lines and the rows represent genes. We estimated an approximate factor embedding 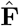 independently in each cell line by solving

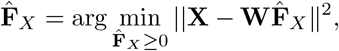

where **W** represents gene loadings of factors from either Love et al (**W**_*L*_) or Dong et al (**W**_*D*_) cohorts and ||.||represents the Frobenius norm. If **V**_*s*_ represents the canonical expression loading matrix obtained from running KSTITCH on sample *s* of the Love et.al or Dong et al cohort, we compute the sample-specific canonical expression projection of **X** as 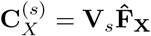 where the first row of 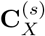 contains CEP1 values of each cell line with subsequent rows containing subsequent CEP scores. The CEP1 scores across cell lines were z-scored separately for each sample-specific loading matrix **V**_*s*_.

In order to compute the overlap of the top-weighted genes in **W**_*L*_ and **W**_*D*_ with known di!erentiation signatures, we downloaded signatures from Tsoi et al [32] which consists of genes up-regulated in melanoma cell lines as melanocytic, transitory,undi!erentiated or neural crest-like, along with intermediate combinations transitory-melanocytic, transitory-neural crest-like and undi!erentiated-neural crest-like. We additionally downloaded MES and MEL signatures consisting of genes up-regulated in MES and MEL cell lines from the Andrews. et al study [36].

### Data and Code Availability

The `kstitch` software package is publicly available at https://github.com/vishakagopalan/kstitch. All code used for the manuscript analyses, the primary analysis notebook, analysis-ready input data, author-generated intermediate files, supplementary table inputs, software environment records, file checksums, and instructions for reproducing the analyses via a Docker container are archived in Zenodo at https://doi.org/10.5281/zenodo.22088660.

## Appendix

### Algorithm 1

Embedding X in preshape space *S*^p^

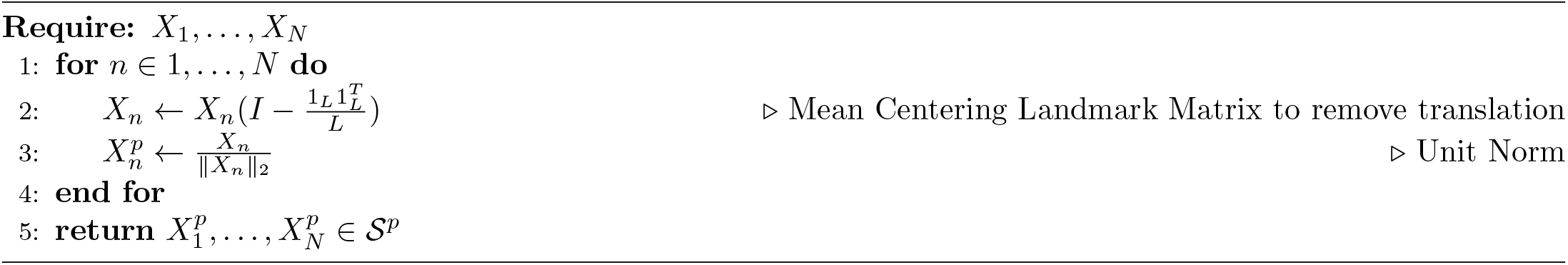

### Algorithm 2

OPA (Orthogonal Procrustes alignment with cyclic shifts)

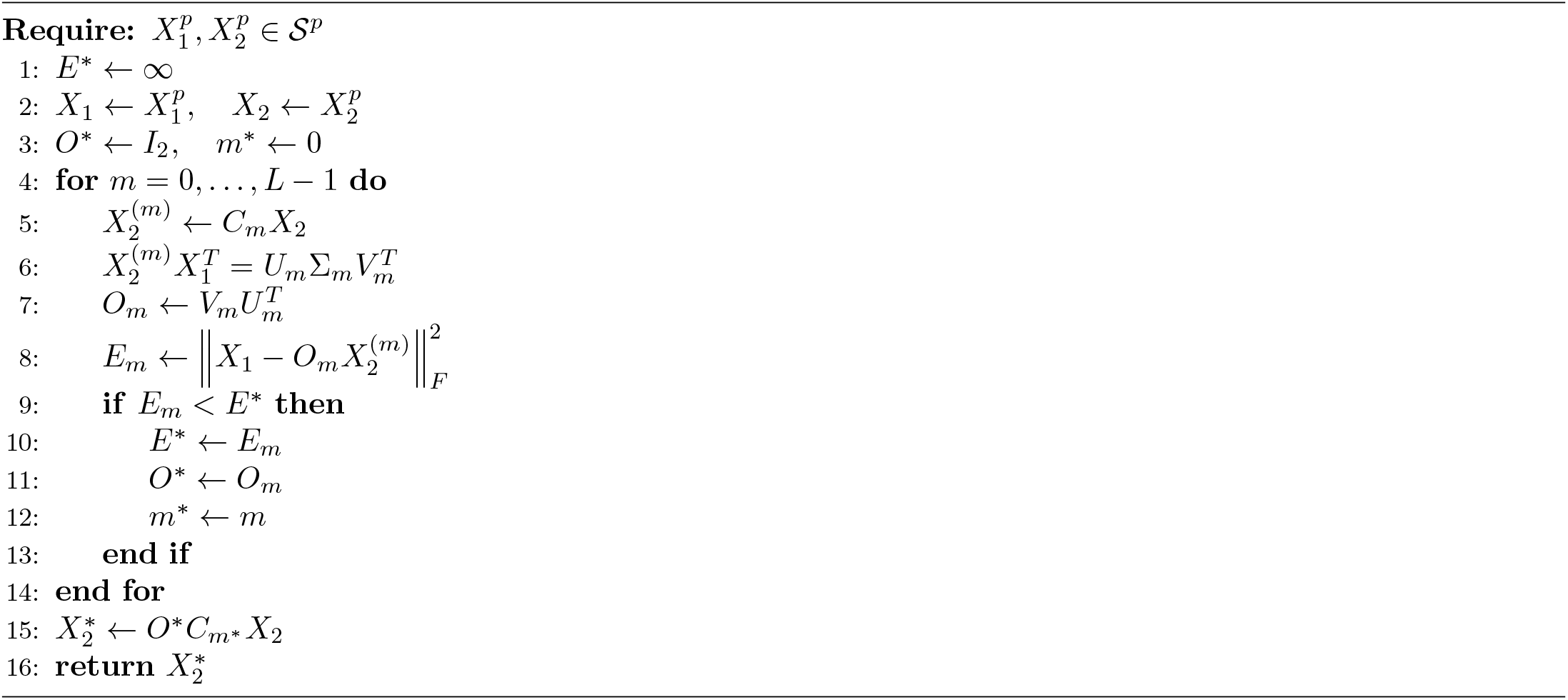

### Algorithm 3

Frechet Mean

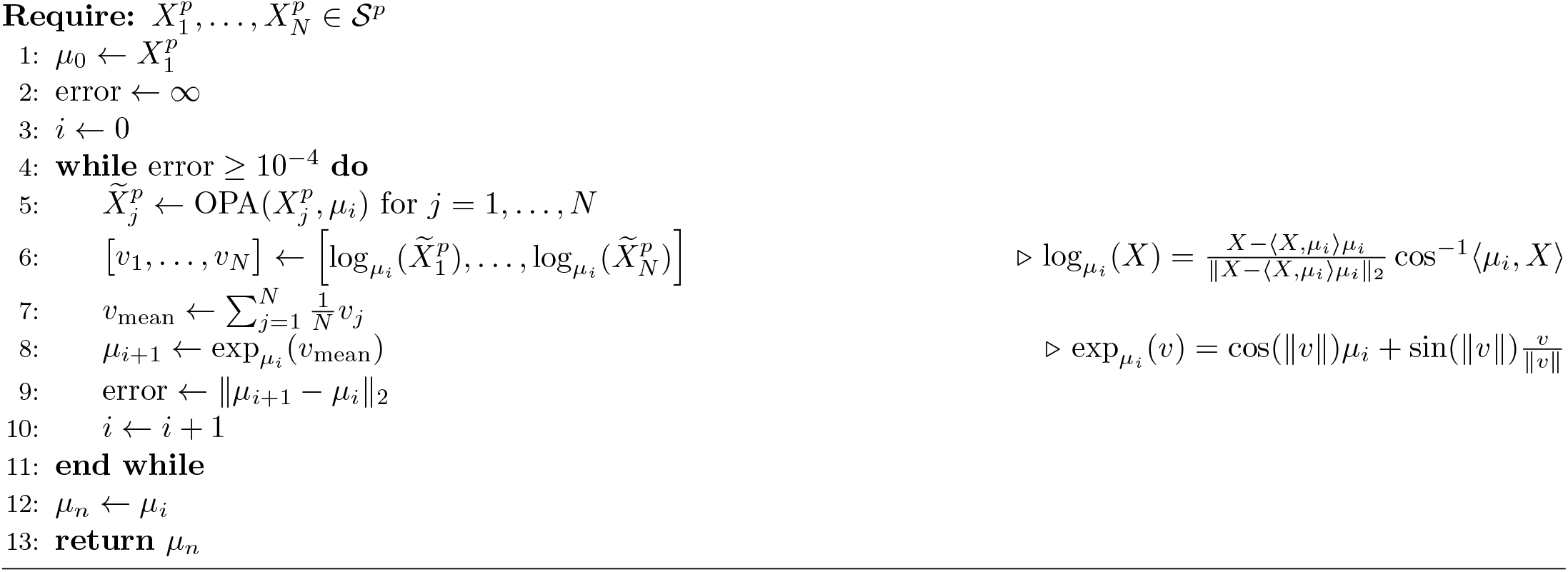

### Algorithm 4

Tangent PCA

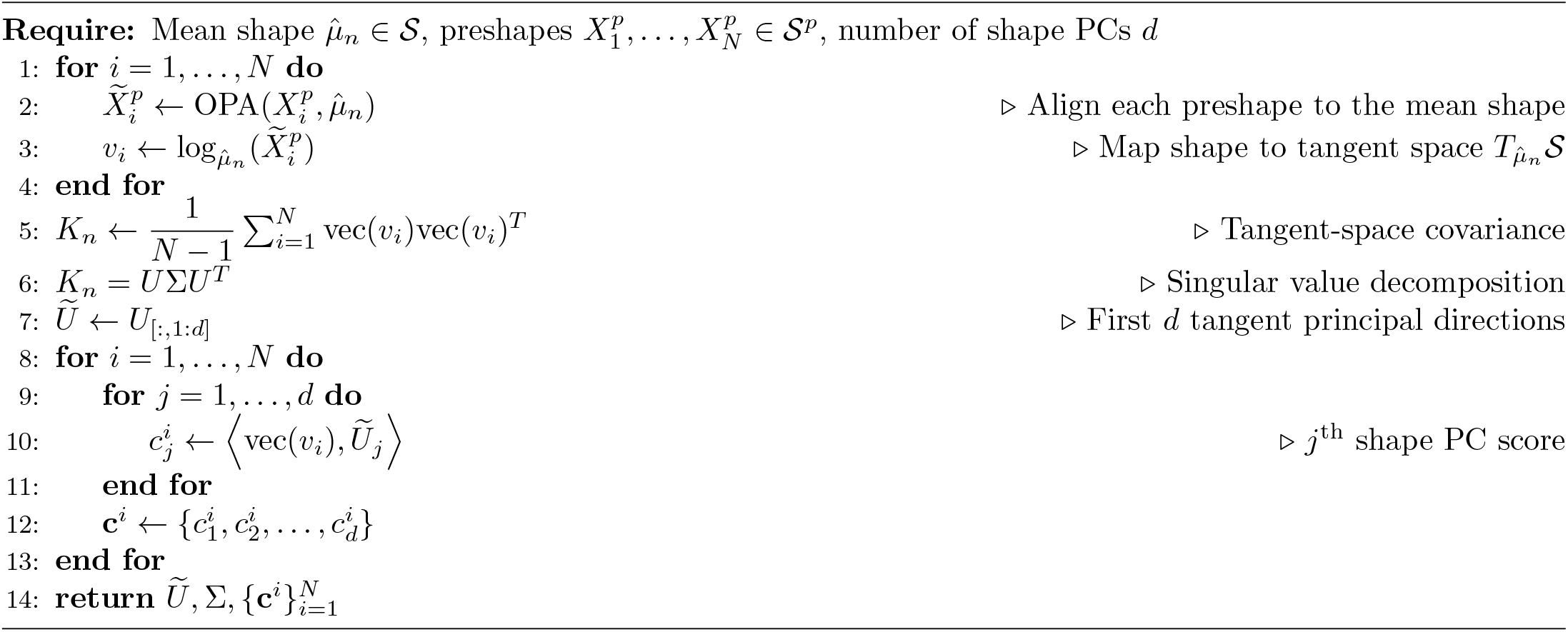

---

**Supp. Figure 1:**
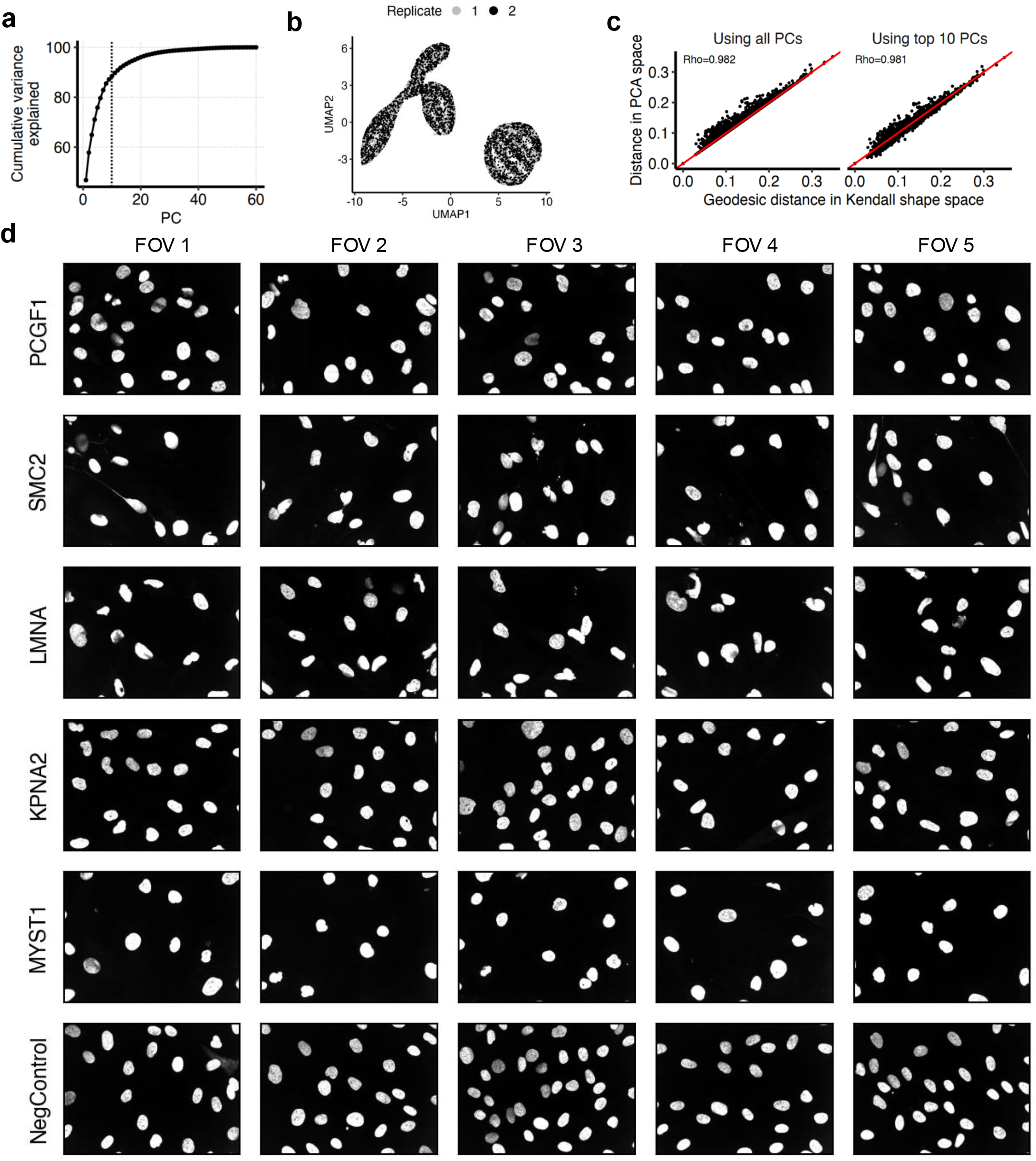
(a) Variance in Schibler. et al data (y-axis) cumulatively explained by considering successive principal components (x-axis). Vertical line drawn at x = 10 represents the number of PCs used in analyses. (b) UMAP embedding of 20,000 sampled nuclei (~1% of dataset), with each point representing a nucleus and colored based on the replicate in which it was imaged. UMAP coordinates were computed based on the first ten shape PCs. (c) Scatter plot of geodesic distances (x-axis) and distances in TPCA space (y-axis) between pairs of nuclei sampled (N = 1000) from Schibler et. al dataset. Distances are calculated based on either all PCs (left panel) or top ten PCs (right panel. (d) Contrast-enhanced images of nuclei (DAPI stains) after knockdown of genes labelled as shape genes in Schibler et. al (PCGF1, SMC2, LMNA) and knockdowns of genes detected by our TPCA approach as shape genes (KPNA2,MYST1) but not detected by Schibler et. al. Nuclei from negative controls (NegControl) are along the bottom row. Each column shows nuclei from different fields of view associated with each perturbation.

**Supp. Figure 2:**
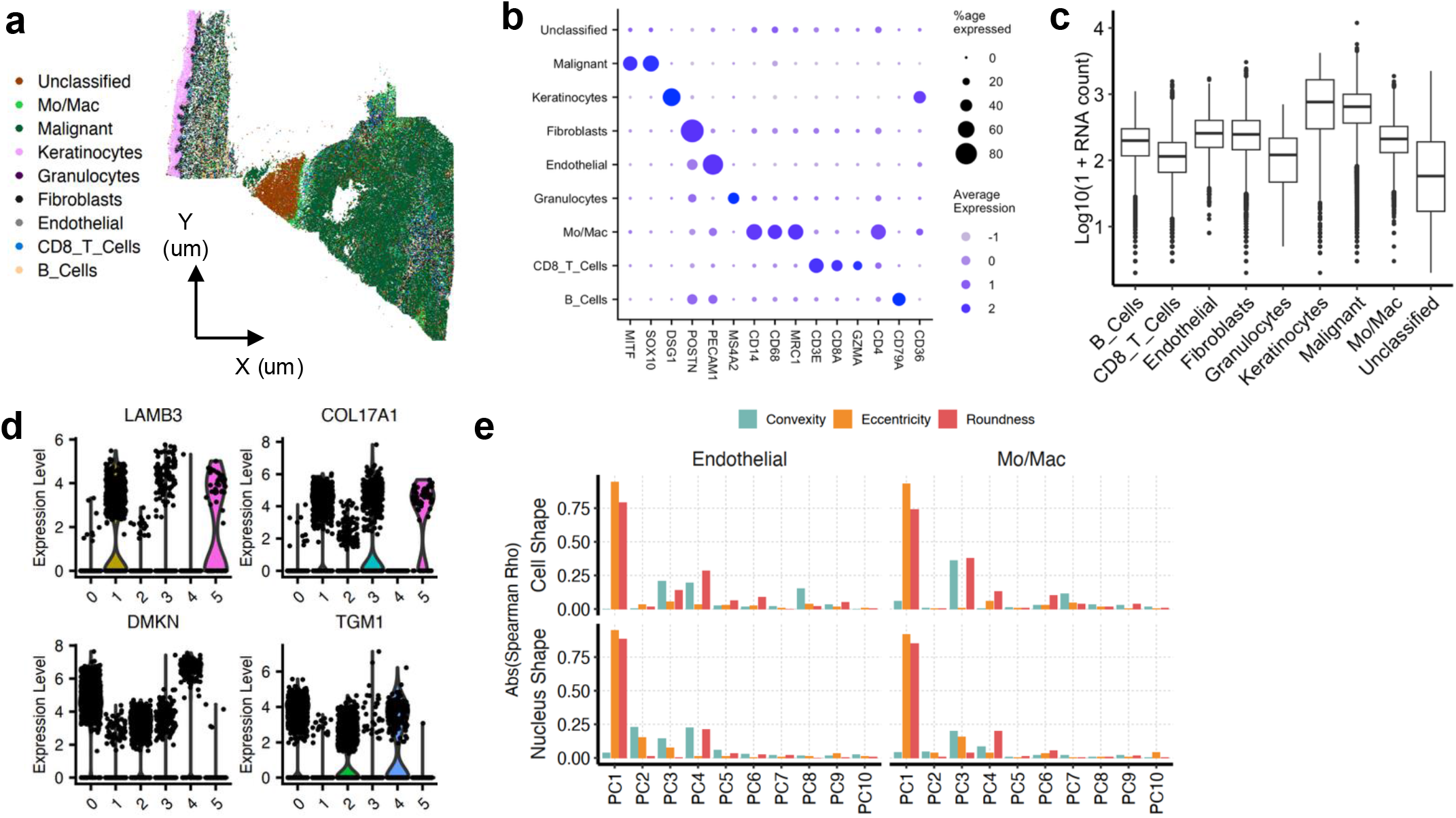
(a) Spatial distribution of annotated cell types in Xenium Prime dermal melanoma dataset. (b) Marker genes used to determine identities of all cell types in Xenium Prime dermal melanoma dataset. (c) Distributions of RNA counts in each cluster. (d) Genes used to label keratinocyte subtypes in dataset. (e) Absolute correlation coefficient (Spearman Rho) between cellular shape PCs and convexity, eccentricity and roundness (top row) of endothelial cells and monocytes/macrophages. Correlation coefficients of nuclear shape PCs with the same shape descriptors are shown in the bottom row.

**Supp. Figure 3:**
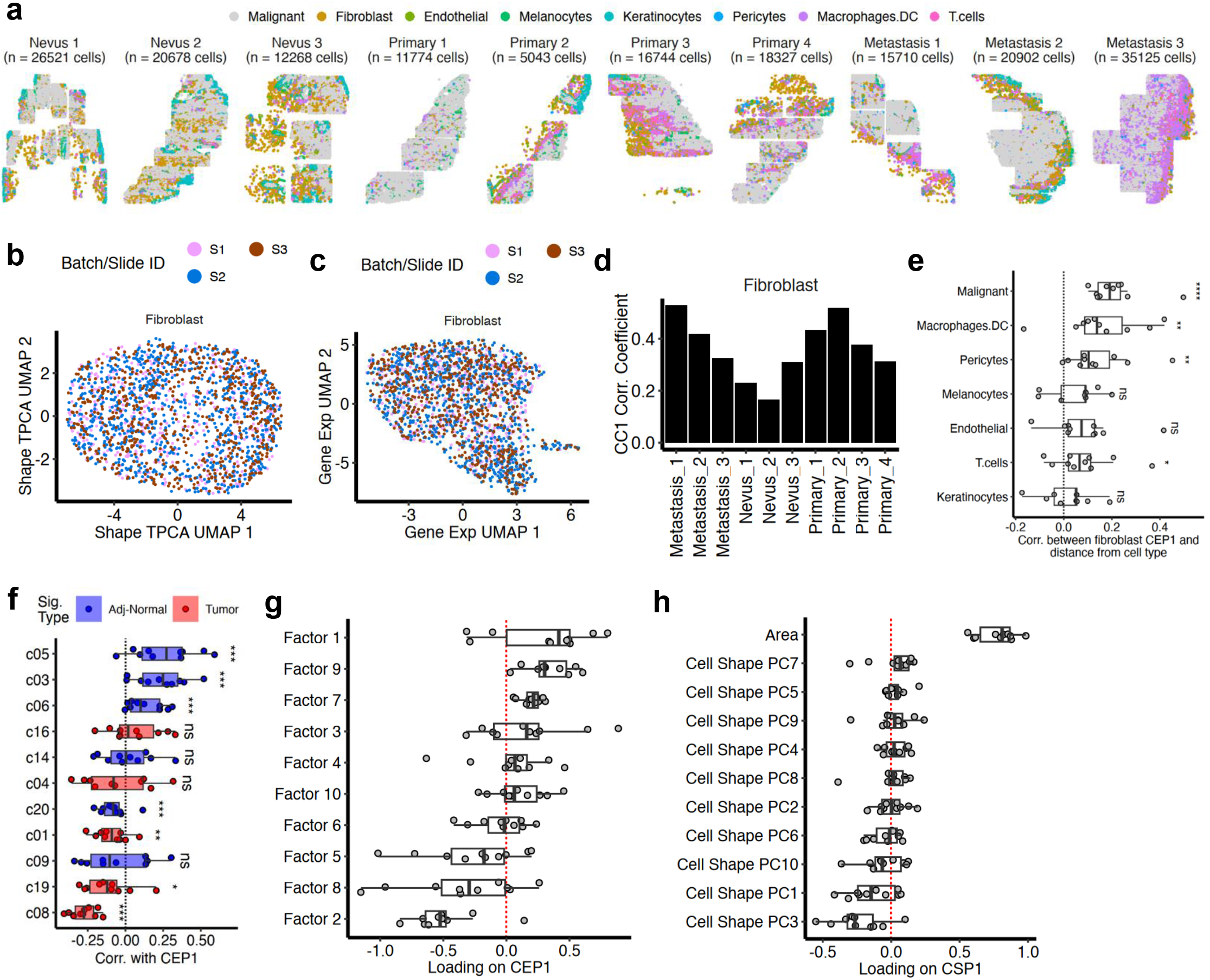
(a) Spatial coordinate plot of all CosMx samples Love et. al melanoma cohort, with color indicating cell type. (b) UMAP coordinates of fibroblasts across all samples based on their cellular shape TPCA embeddings and colored by the CosMx slide from which they originate. (c) UMAP coordinates computed from NMF embeddings of fibroblasts pooled across samples. (d) Correlation coefficient associated with the first canonical correlation (between CSP1 and CEP1) when STITCH is run separately on fibroblasts from each sample of Love et. al cohort. Bars in black indicate significant (p < 0.05) correlations in that sample. (e) Correlation of fibroblast CEP1 score from each sample with distance from each cell type. Each dot represents a different sample. (f) Correlation between fibroblast CEP1 of each sample and expression of fibroblast signatures from Gao et. al. Each dot represents correlations from a single sample. (g) Loadings of each shape PC and cellular area on CSP1 from fibroblasts of each sample. Each dot represents loadings from a single sample. (h) Loadings of each factor on CEP1 from fibroblasts of each sample. Each dot represents loadings from a single sample.

**Supp. Figure 4:**
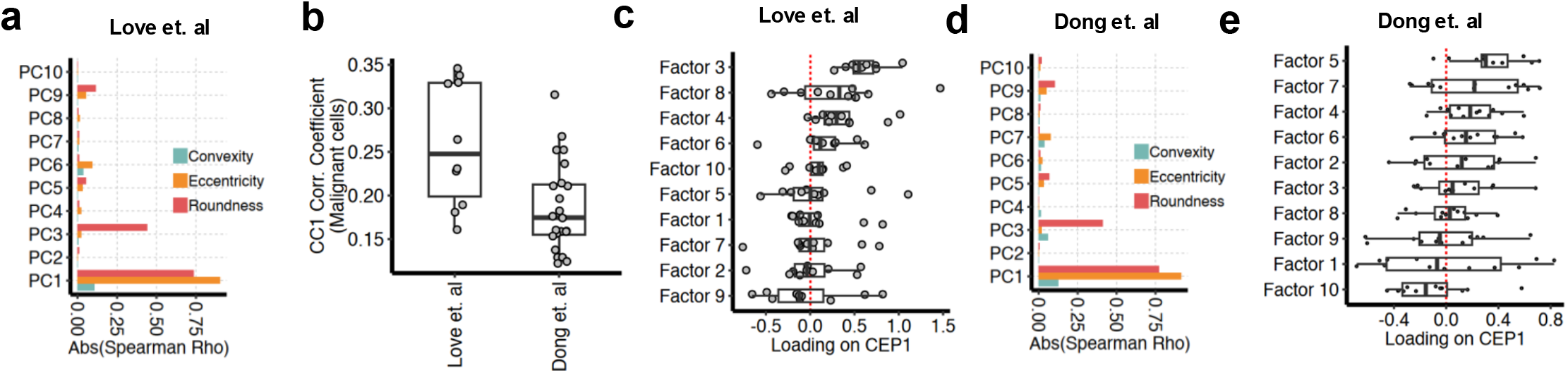
(a) Correlation of malignant cell boundary PC shape modes from Love et. al cohort with cell eccentricity, roundness and convexity. (b) Distribution of CC1 correlation coefficients obtained from running STITCH on malignant cells from samples Love et. al cohort (n=10) and Dong et. al cohort (n=25). (c) Loadings of gene expression factors derived from malignant cells in Love et. al cohort on CEP1. Each dot represents a loading in a different sample from Love et. al cohort. (d) Correlation of malignant cell boundary PC shape modes from Dong et. al cohort with cell eccentricity, roundness and convexity. (e) Loadings of gene expression factors derived from malignant cells in Dong et. al cohort on CEP1. Each dot represents a loading in a different sample from Dong et. al cohort.

